# Bitter Taste Receptor Agonists Induce Apoptosis in Papillary Thyroid Cancer

**DOI:** 10.1101/2024.10.18.618693

**Authors:** Kimberly Wei, Brianna L. Hill, Zoey A. Miller, Arielle Mueller, Joel C. Thompson, Robert J. Lee, Ryan M. Carey

**Author notes:** Corresponding Author: Ryan M. Carey, MD, Department of Otorhinolaryngology – Head and Neck Surgery Hospital of the University of Pennsylvania, 3400 Spruce Street, 5^th^ floor Ravdin Building Philadelphia, PA 19104.

## Abstract

**Background:** Papillary thyroid carcinoma (PTC) is the most common thyroid malignancy, with a 20% recurrence rate. Bitter taste receptors (T2Rs) and their genes (*TAS2Rs*) may regulate survival in solid tumors. This study examined T2R expression and function in PTC cells.

**Methods:** Three PTC cell lines (MDA-T32, MDA-T68, MDA-T85) were analyzed for expression using RT-qPCR and immunofluorescence. Live cell imaging measured calcium responses to six bitter agonists. Viability and apoptosis effects were assessed using crystal violet and caspase 3/7 activation assays. Genome analysis of survival was conducted.

**Results:** *TAS2R14* was consistently highly expressed in all cell lines. Five bitter agonists produced significant calcium responses across all cell lines. All bitter agonists significantly decreased viability and induced apoptosis. Higher *TAS2R14* expression correlated with better progression-free survival in patients (p<0.05).

**Conclusions:** T2R activation by bitter agonists induces apoptosis and higher *TAS2R* expression is associated with survival, suggesting potential therapeutic relevance in thyroid cancer management.

## Introduction

Papillary thyroid cancer (PTC) is a differentiated epithelial tumor that arises from the follicular cells of the thyroid. It is the most common thyroid neoplasm, constituting about 80% of all cases and its incidence and prevalence are on the rise. Risk factors include female gender (3:1 female to male ratio), previous exposure to ionizing radiation, and rare hereditary conditions such as Cowden syndrome.^1^ In general, PTC has the best overall prognosis out of all the thyroid carcinomas with a 93% 10-year survival rate.^2^ However, recurrence rates range from 15-30% due to PTC’s propensity for metastasis through the lymphatics. Patients with identified lymph node (LN) metastases have worse relapse-free and overall survival.^3–5^ In one large scale study, the recurrence of PTC presenting with LN metastases was reported to be 31.5% and out of the patients with recurrent PTC, 40% died of thyroid cancer.^6^

The current standard of care is based on the extent of disease and includes surgery (thyroid lobectomy or total thyroidectomy) potentially followed by radioactive iodine (RAI) ablation and life-long thyroxine therapy.^7^ Recurrences are managed either through surgery RAI, external beam radiation therapy, observation, or percutaneous interventions such as ethanol injection.^8^ However, there is still a need for alternative therapies, especially in metastatic PTC. On the other spectrum, very indolent PTC with very low risk of metastasis is often at risk of overtreatment. Active surveillance is becoming more common in managing this type of PTC, but other minimally invasive interventions could improve the management of low-risk PTC.^9^

Bitter taste receptors (taste family 2 receptors or T2Rs) and their associated genes (*TAS2Rs*) have recently been studied in a variety of solid tumors including breast, ovarian, and head and neck squamous cell carcinoma.^10–14^ T2Rs are G protein-coupled receptors (GPCRs) that signal through Gα to decrease cyclic adenosine monophosphate (cAMP) and Gβγ to activate phospholipase C and calcium (Ca^2+^) release from the endoplasmic reticulum (ER).^15–19^ There are 25 human T2R proteins and genes.^20^ Initially identified on the tongue for their role in taste, T2Rs have been identified in many other malignant and non-malignant tissues. Physiologically, T2Rs are involved in various normal chemosensory functions including serving as sentinels for innate immunity against bacteria.^21–23^ In cancers, these receptors have been found to have altered expression compared to normal tissues^24,25^ and play a role in regulating multidrug resistance transporters, as well as migration and anticancer functions such as apoptosis.^26–29^

While there have been prior studies on T2Rs in solid tumors^11,12,25^, there has yet to be any study that explores the presence and function of T2Rs in PTC. The aim of this project is to characterize the expression and function of T2Rs in response to bitter agonists in PTC cells with the goal of generating potential therapeutic roles in the future. We hypothesized that T2Rs would regulate viability as previously described in head and neck squamous cell carcinoma.^11,12^ Our study demonstrates that T2Rs are expressed in PTC cells and can be activated to induce apoptosis.

## Materials and Methods

This study was deemed exempt from review by the Institutional Review Board (IRB) of the University of Pennsylvania Health System (2023-HSF1203).

### Cell Culture

MDA-T32, MDA-T68, and MDA-T85 cell lines were obtained from ATCC (Manassas, VA, USA). MDA-T32 (CRL-3351) is a well-differentiated papillary carcinoma. MDA-T68 (CRL-3353) is a papillary carcinoma of the follicular variant. MDA-T85 (CRL-3354) is a metastatic papillary carcinoma from a lymph node. All cell lines were grown in submersion in high glucose Dulbecco’s modified Eagle’s medium (Corning; Glendale, AZ, USA) with 10% FBS (Genesee Scientific; El Cajon, USA), and penicillin/streptomycin mix (Gibco; Gaithersburg, MD, USA).

### Reverse transcription quantitative PCR (RT-qPCR)

Cell cultures were resuspended in TRIzol (Thermo Fisher Scientific; Waltham, MA, USA). RNA was isolated and purified (Direct-zol RNA kit; Zymo Research), reverse transcribed via High-Capacity cDNA Reverse Transcription Kit (Thermo Fisher Scientific), and quantified using TaqMan qPCR probes for *TAS2R1, TAS2R3, TAS2R4, TAS2R5, TAS2R7, TAS2R8, TAS2R9, TAS2R10, TAS2R13, TAS2R14, TAS2R16, TAS2R19, TAS2R20, TAS2R38, TAS2R39, TAS2R40, TAS2R41, TAS2R42, TAS2R43, TAS2R44, TAS2R45, TAS2R46, TAS2R47, TAS2R50, TAS2R60,* and *UBC* (QuantStudio 5; Thermo Fisher Scientific). UBC was used as an endogenous control due to its stable expression in cancer cells.^30,31^

### Live Cell Imaging

For calcium Ca^2+^ imaging, cells were loaded with 5µM of Fluo-4-AM (Thermo Fisher Scientific) for 60 min at room temperature in the dark. Hank’s Balanced Salt Solution (HBSS) buffered with 20mM HEPES (pH 7.4) was used as an imaging buffer, containing 1.8mM Ca^2+^. Cells were imaged in 48-well plastic plates using a Nikon Eclipse TS100 (20x 0.75 NA PlanApo objective), FITC filters (Chroma Technologies), QImaging Retiga R1 camera (Teledyne; Tucson, AZ, USA), MicroManager^32^, and Xcite 120 LED Boost (Excelitas Technologies; Mississauga, Canada). Bitter agonists were dissolved in HBSS. At the start of the experiment, 150uL of bitter agonist in HBSS was added to 150uL of HBSS.

T2R14 inhibitor LF1 was dissolved in DMSO at 500mM and used at 1:1000 as indicated.^33^ Stock solutions were stored at -20°C. Experimental wells were incubated with HBSS and LF1 and experiments were conducted by adding 150uL of dissolved bitter agonist onto cells submerged in 150uL of HBSS and LF1.

### Cell Viability

Cells were incubated in phenol-free high glucose Dulbecco’s modified Eagle’s medium (Corning; Glendale, AZ, USA) with 10% FBS (Genesee Scientific; El Cajon, USA), and penicillin/streptomycin mix (Gibco; Gaithersburg, MD, USA) with bitter agonists for 24 hours at 37°C. Crystal violet (0.1% in deionized water with 10% acetic acid) was used to stain remaining adherent cells. Stains were washed with deionized water and left to dry at room temperature.

Stains were dissolved with 30% acetic acid in deionized water. Absorbance values at 590nm were measured on a Tecan (Spark 10M; Mannedorf, Switzerland).

### Apoptosis Measurements

CellEvent Caspase 3/7 dye was added to cells in combination with bitter agonists.

Fluorescence was measured at 495nm excitation and 540nm emission. Live cell images were taken on Olympus IX-83 microscope (20x 0.75 NA PlanApo objective), FITC filter (Chroma Technologies), Orca Flash 4.0 sCMOS camera (Hamamatsu, Tokyo, Japan), Meta-Fluor (Molecular Devices, Sunnyvale, CA USA), and XCite 120 LED Boost (Excelitas Technologies). Fluorescence values were quantified through image analysis on Image J.

### Immunofluorescence

Cells were fixed on glass with 4% formaldehyde in PBS w/ Ca^2+^ and Mg^2+^. Cells were blocked and permeabilized in blocking buffer containing PBS with Ca^2+^ and Mg^2+^, 3% normal donkey serum, 1% BSA, 0.2% saponin, 0.1% Triton X-100. Cells were incubated overnight in the primary antibody (Anti-TAS2R14, Cat# C386266; RRID: <u>AB_1936354</u>) at 1:100. Cells were then incubated in secondary antibody (Anti-rabbit Alexa Fluor 647, Cat# A31537; RRID: <u>AB_2536183</u>) at 1:500 for 2 hours. Phalloidin was used to stain actin at 1:400 and DAPI was used to stain the nucleus. Images were taken on Olympus Live Cell Imaging System (system as described in apoptosis measurements) with 60x oil objective.

### Western Blotting

Protein was harvested using lysis buffer (10 mM Tris pH 7.5, 150 mM NaCl, 10 mM KCl, 0.5% deoxycholate, 0.5% Tween, 0.5% IGePawl, 0.1% SDS). DC Protein Assay (BioRad; Hercules, CA, USA) was used to quantify protein content. 50µg of protein were loaded into gel (Bis-Tris 4-12%, 1.5 mm) with 4x loading buffer (200 mM Tris pH 6.8, 40% glycerol, 8% SDS, 0.1% Bromophenol Blue), and 5% b-mercaptoethanol. Gel was run using MES running buffer (Tris Base 50 mM pH 7.3, MES 50 mM, SDS 0.1%, EDTA 1 mM). Novex Sharp Pre-Stained Protein Standard, SeeBlue Plus2 Pre-stained Protein Standard (Thermo Fisher), or Odyssey One-Color Protein Molecular Weight Marker (LI-COR) were used as molecular markers. Gel was transferred using Bis-Tris transfer buffer (25 mM bicine, 25 mM bis-tris, 1 mM EDTA, 10% methanol). Blots were blocked for 1 hour in TBST (24.8 mM tris acid & base, 1.5 NaCl, 0.5% Tween-20) with 5% milk. Primary antibodies (Anti-TAS2R14, Cat# C386266; RRID: <u>AB_1936354</u>) were used 1:1000 or 1:500 with HRP-conjugated chemiluminescent secondary antibodies (Anti-rabbit IgG, HRP Conjugated, Cat# SC-2004; RRID: <u>AB_631746</u>) 1:1000 in TBST with 5% BSA (primary) or milk (secondary). Blots were imaged using Clarity Max Western ECL Substrate (BioRad) using ChemDoc MP Imaging System (BioRad).

### Analysis of The Cancer Genome Atlas (TCGA) Thyroid Carcinoma (THCA) Cohort

RSEM expected, Deseq2 standardized gene expression counts for *TAS2R4*, *TAS2R10*, and *TAS2R14* were obtained from the THCA TCGA TARGET GTEx study for both primary tumor (n=504) and normal tissue (n=338) using the University of California Santa Cruz (UCSC) Xena Browser (https://xenabrowser.net/).^34^ Gene expression values were stratified by sample type (primary tumor or normal) and plotted in GraphPad Prism (San Diego, CA, USA). Overall and progression-free survival analyses were performed in the UCSC Xena Browser using the RSEM expected, Deseq2 standardized counts for *TAS2R4*, *TAS2R10*, and *TAS2R14*.

### Statistical Analysis

Data was analyzed in GraphPad Prism (San Diego, CA, USA). t-tests (two comparisons only) or one-way ANOVA (more than two comparisons) were conducted. Both paired and unpaired tests were used when appropriate. Dunnett’s posttests for one-way ANOVA were used when needed.

## Results

### T2Rs are expressed in PTC cells

The expression of all 25 human *TAS2Rs* was analyzed in three human PTC cell lines (MDA-T32, MDA-T68, MDA-T85) by reverse transcription quantitative PCR (RT-qPCR) (**Fig. 1A**).

**Figure 1.**
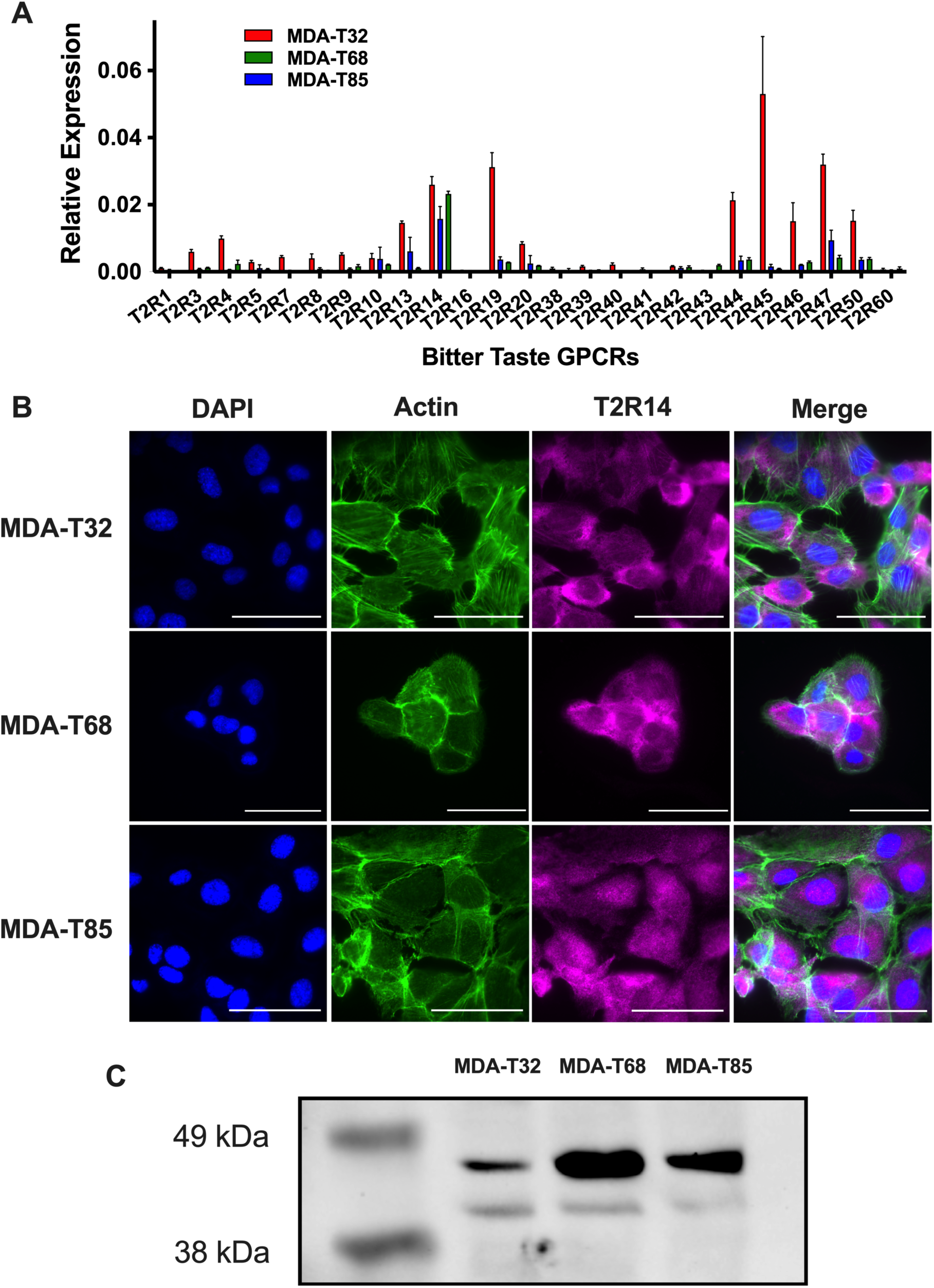
Bitter (T2R) taste receptors are expressed in PTC cells. **A)** Quantitative PCR (qPCR) expression analysis of *TAS2R* transcripts in three PTC cell lines. Expression of each *TAS2R* mRNA was normalized to ubiquitin C (UBC) housekeeping gene. **B)** T2R14 protein expression in PTC cell lines (MDA-T32, MDA-T68, MDA-T85) via immunofluorescence stain with DAPI (nucleus) and phalloidin (actin). Scale bar represents 130μm. Each antibody was compared to secondary only control at the same microscope settings. **C)** T2R14 protein expression in MDA-T32, MDA-T68, and MDA-T85 cells via western blot (1:500 primary antibody).

Variable expression was demonstrated with cell line MDA-T32 showing the highest expression levels across several *TAS2Rs* compared to MDA-T68 and MDA-T85. *TAS2R14* expression was consistently one of the highest in all three cell lines, with an expression level of around 2% relative to housekeeping gene UBC. Therefore, confocal immunofluorescence microscopy was used to visualize T2R14 localization. Fixed MDA-T32, MDA-T68, and MDA-T85 cells were stained with antibodies targeting T2R14. T2R14 was found to be expressed, likely in the endoplasmic reticulum (ER), with a possibility of also being on the extracellular plasma membrane (**Fig. 1B**). To further validate expression of T2R14, a western blot was performed, and expression was confirmed in both MDA-T32 and MDA-T68 cells (**Fig. 1C**).

### Bitter agonist stimulation of bitter taste receptors induces intracellular Ca^2+^ responses in PTC cells

To determine whether T2Rs in PTC cells are functional, bitter agonist-induced intracellular Ca^2+^ changes in MDA-T32, MDA-T68, and MDA-T85 were examined. Ca^2+^ responses were recorded in living PTC cells loaded with Fluo-4.^35^ Six different bitter agonists were used: denatonium, diphenhydramine, flufenamic acid (FFA), quinine, thujone, and lidocaine. Each target a specific subset of T2Rs (**Table 1**). In MDA-T32 and MDA-T68 cells, all agonists induced statistically significant Ca^2+^ responses. In MDA-T85 cells, all except thujone produced significant Ca^2+^ responses (**Fig. 2A-B**).

**Figure 2.**
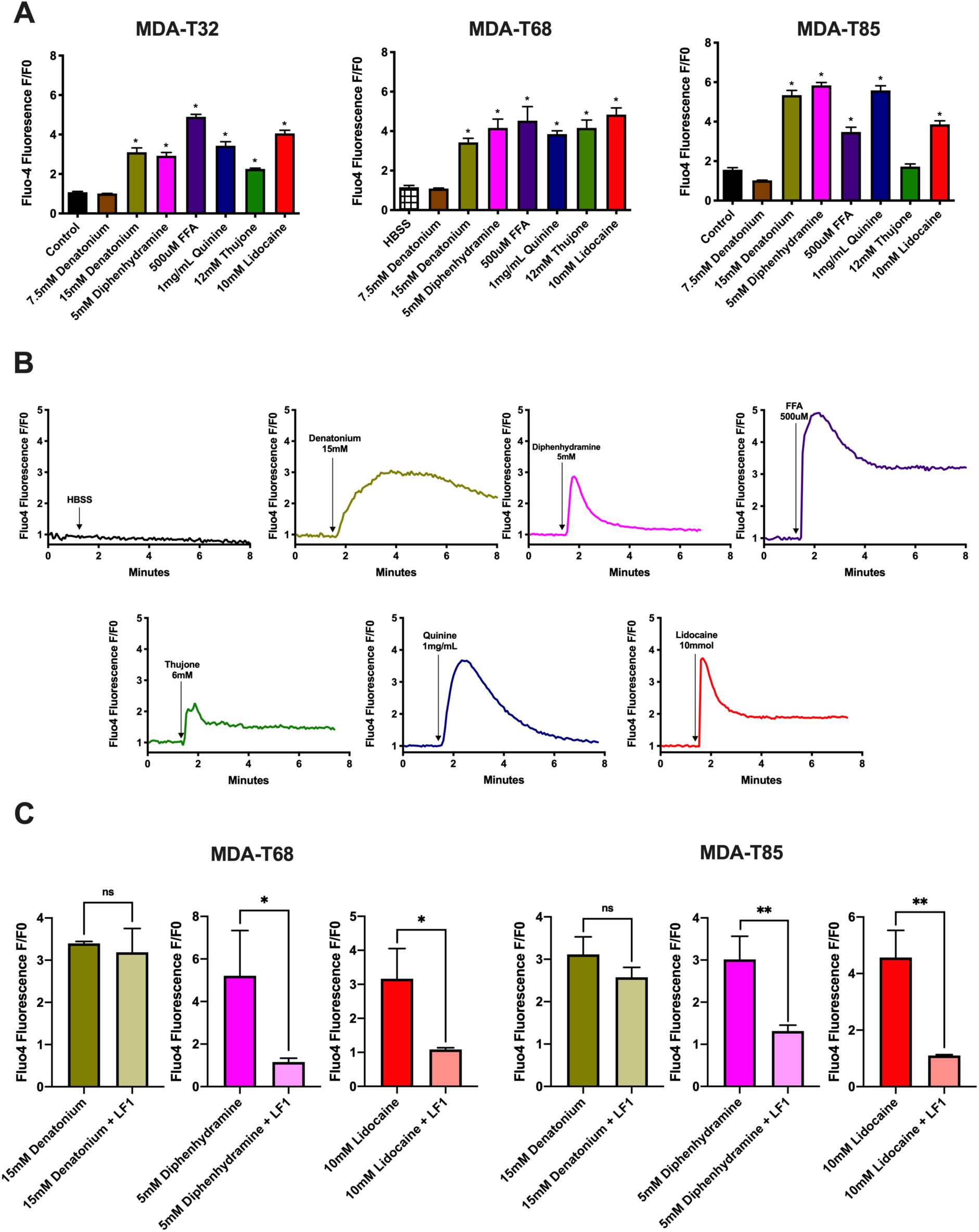
Bitter agonists induce Ca^2+^ responses via T2Rs in PTC cells. **A)** Peak Ca^2+^ responses to bitter agonists in three PTC cell lines; mean ± SEM with >3 experiments using separate cultures. Significance by one-way ANOVA with Bonferroni post-test comparing HBSS with each bitter agonist. **B)** MDA-T32 Ca^2+^ response over time. **C)** MDA-T68 and MDA-T85 peak Ca^2+^ responses with denatonium, diphenhydramine, and lidocaine ± 1 hour before incubation with T2R14 inhibitor LF1. *p < 0.05; **p < 0.001; ns, no statistical significance (or no indication).

**Table 1.** Bitter agonists and respective activated T2Rs^69^.

| **Bitter Agonist** | **Activated T2Rs** |
| --- | --- |
| Denatonium benzoate | T2R4, T2R8, T2R10, T2R13, T2R39, T2R43,<br>T2R46, T2R47 |
| Diphenhydramine | T2R14, T2R40 |
| Flufenamic Acid (FFA) | T2R14 |
| Quinine | T2R4, T2R7, T2R10, T2R14, T2R39, T2R40,<br>T2R43, T2R44, T2R46 |
| Thujone | T2R10, T2R14 |
| Lidocaine | T2R14 |

To further test if intracellular Ca^2+^ changes were due to T2R activation, MDA-T68 and MDA-T85 cells were incubated with LF1, a T2R14 inhibitor.^36^ Three bitter taste agonists were used. Both lidocaine and diphenhydramine activate T2R14, while denatonium was used as a positive control because it does not activate T2R14. In both cell lines, there was a significant decrease in Ca^2+^ response in cells treated with LF1 prior to lidocaine and diphenhydramine, but there was no change in response to denatonium (**Fig. 2C**). These findings suggest that bitter taste agonists induce Ca^2+^responses through T2R14 bitter taste receptors.

### Bitter agonists decrease cell viability and induce apoptosis in PTC cells

To examine the effects of bitter taste receptor activation by bitter agonists, a crystal violet viability assay was conducted in all three cell lines. After 24 hours, all bitter agonists significantly decreased cellular viability in MDA-T32 cells. In MDA-T68 and MDA-T85 cells, all agonists significantly decreased viability except for FFA (**Fig. 3**).

**Figure 3.**
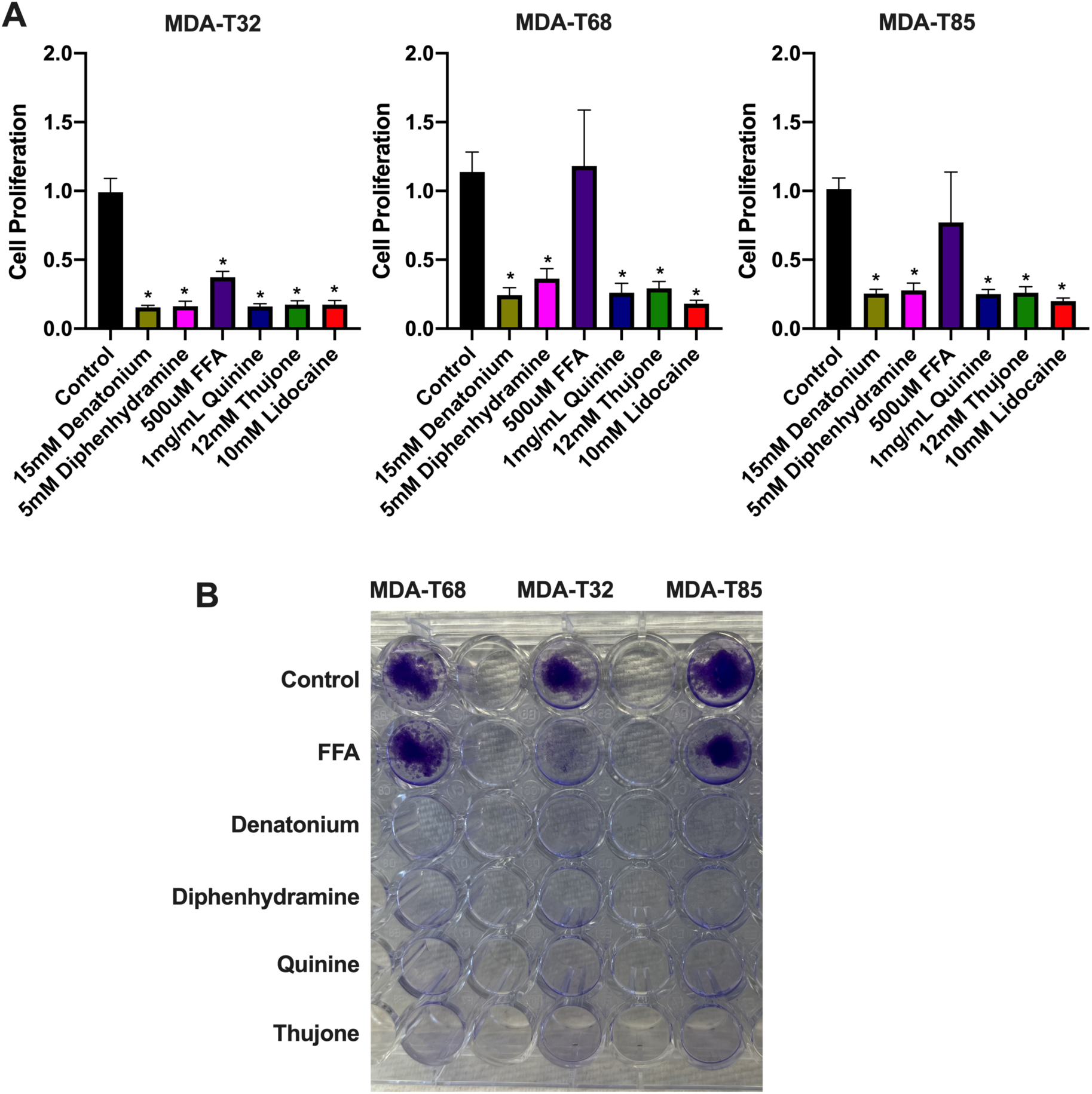
Bitter agonists inhibit cell viability in PTC cells. **A)** MDA-T32, MDA-T68, and MDA-T85 viability response to bitter agonists via crystal violet assay. **B)** Representative image of crystal violet assay. *p < 0.05; ns, no statistical significance (or no indication).

To determine if the effects on viability are due to bitter agonists being pro-apoptotic, a CellEvent assay that measures caspase-3 and -7 cleavage in PTC cells was conducted. The CellEvent dye fluoresces upon caspase cleavage, indicating apoptosis.^37–39^ Diphenhydramine, thujone, and lidocaine were found to induce significant apoptotic responses in MDA-T32 cells. In MDA-T68 cells, diphenhydramine, FFA, thujone, and lidocaine induce apoptosis, while denatonium, diphenhydramine, thujone, and lidocaine induce apoptosis in MDA-T85 cells (**Fig. 4A-B**). DIC images taken in parallel to the CellEvent assay show abnormal cellular morphology (rounding and lifting) consistent with apoptosis. Apoptotic responses were shown to start as early as 3 hours during a 12-hour time course (**Fig. 4C**). These findings support the anti-viability and pro-apoptotic effects of bitter agonists on PTC cells including compounds with approved clinical applications such as lidocaine.

**Figure 4.**
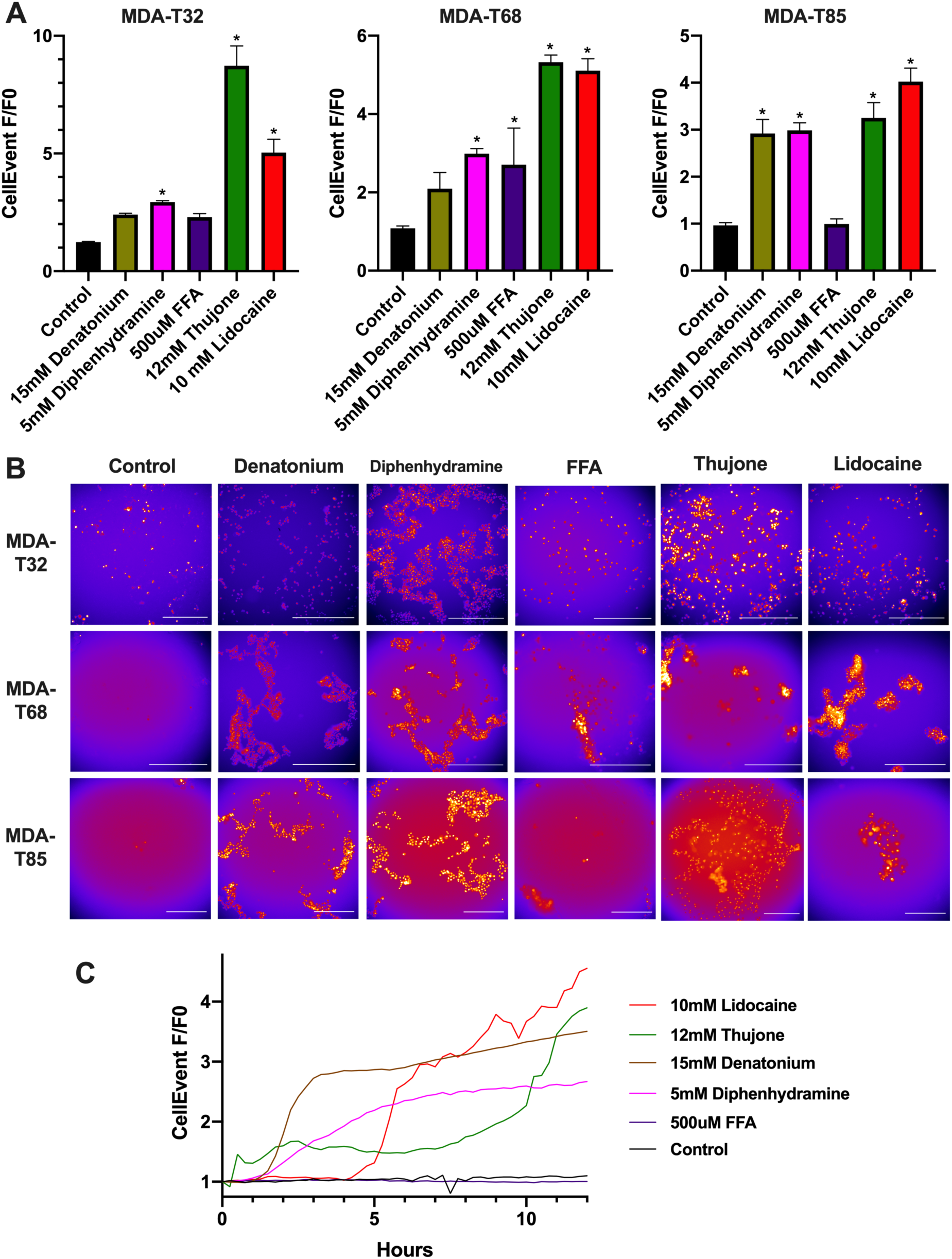
Bitter agonists induce apoptosis in PTC cells. **A)** CellEvent (caspase-3 and -7 indicator) fluorescence at 12 hours (mean ± SEM). Significance determined by one-way ANOVA with Bonferroni post-test between control and bitter agonist treated cells. **B)** Representative images of CellEvent caspase cleavage at 12 hours with bitter agonists. **C)** MDA- T85 representative trace of CellEvent fluorescence over 12 hours with bitter agonists. Scale bar represents 130μm. *p < 0.05; ns, no statistical significance (or no indication).

### Increased T2R14 expression may be associated with better PTC progression free survival

Data above suggest T2Rs activate apoptosis and limit cell viability in PTC cells in vitro, which prompted exploration of the possible in vivo effects of T2Rs. Specifically, we investigated the impact of increased *TAS2R* expression on survival in PTC using TCGA. Analysis included 504 cases of PTC diagnosed between 1993 and 2013 with mRNA expression data available. In concordance with our experimental data, *TAS2R14* was found to have the highest expression amongst the *TAS2R*s in PTC cells (**Fig. 5A**). When analyzing the difference in expression of certain *TAS2R*s in normal vs tumor thyroid cells, *TAS2R4* (FC=-1.08, p<0.0001), *TAS2R10* (FC=-1.24, p<0.0001) and *TAS2R14* (FC=-1.03, p<0.001) were all found to be significantly lower in tumor cells compared to normal thyroid cells (**Fig. 5B**). Kaplan-Meier survival analysis of cases with high vs low *TAS2R14* expression was not significant for overall survival (OS) (p=0.62 by log-rank test, **Fig. 5C**) but did demonstrate significantly improved progression-free survival (PFS) for cases with increased *TAS2R14* expression (p=0.01 by log-rank test; **Fig. 5D**). The 10-year progression-free survival for higher *TAS2R14* expression was 91% vs. 86% for lower *TAS2R14* expression. The OS and PFS for other T2Rs (*TAS2R4* and *TAS2R10*) were analyzed but not found to be significant (**Supp. Fig. 1**).^40^

**Figure 5.**
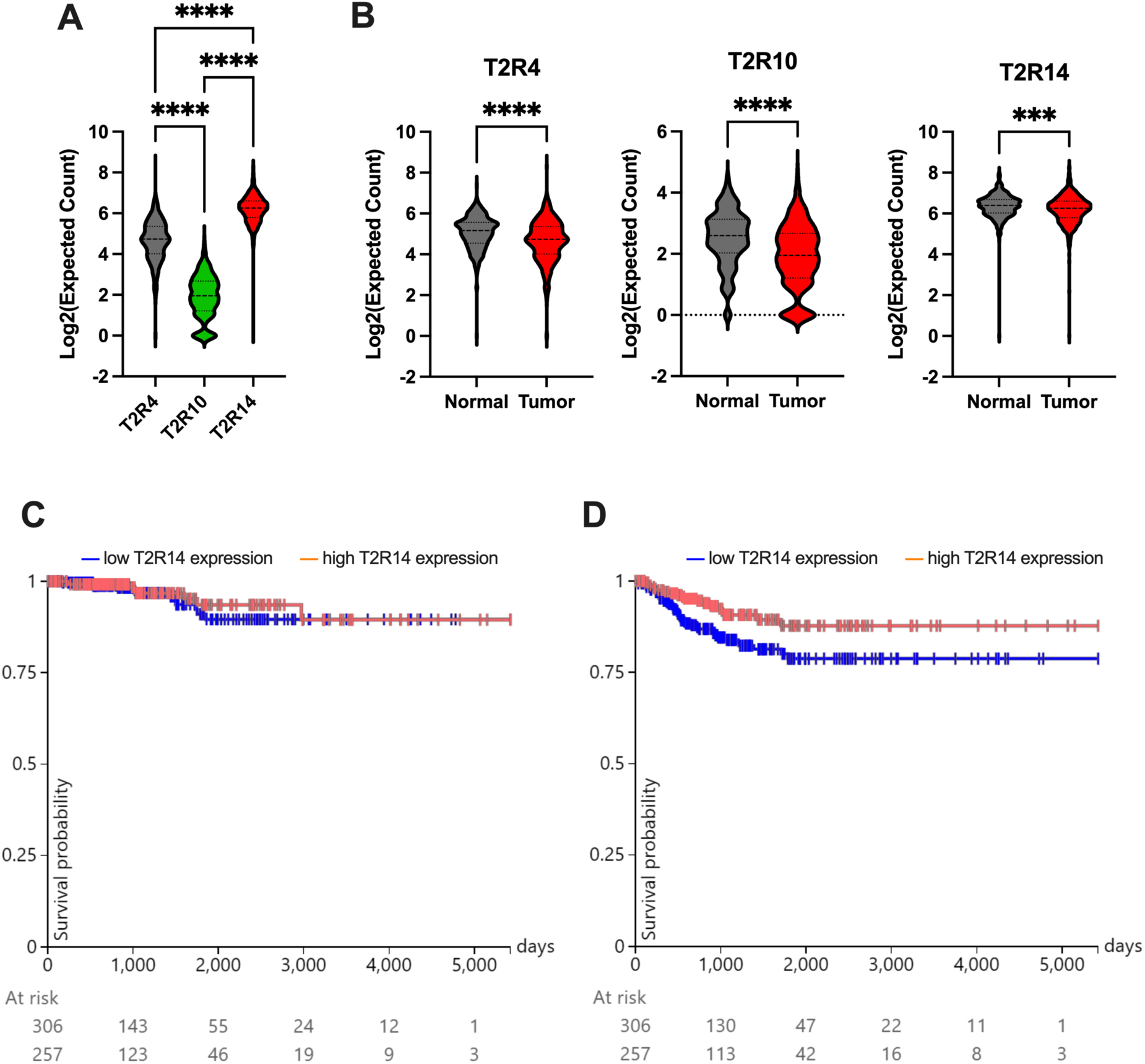
Higher T2R14 expression correlates with improved PFS. **A)** Expression of three *TAS2R*s (*TAS2R4*, *TAS2R10*, *TAS2R14*) in PTC tumors in the Cancer Genome Atlas. Significance by one-way ANOVA. For TAS2R4 vs. TAS2R10: μ*_TAS2R4_*=4.66, μ*_TAS2R10_*=1.92, μd=2.74, σd=0.059. For TAS2R4 vs. TAS2R14: μ*_TAS2R4_*=4.66, μ*_TAS2R14_*=6.19, μd=-1.53, αd=0.059. For TAS2R10 vs. TAS2R14: μ*_TAS2R10_*=1.92, μ*_TAS2R14_*=6.19, μd=-4.27, σd=0.059. ****p<0.0001 **B)** Expression of three *TAS2R*s (*TAS2R4*, *TAS2R10*, *TAS2R14*) in papillary thyroid tumor tissue compared to normal thyroid tissue. Significance by T-test. For *TAS2R4*: μ_normal_=5.06, μ_tumor_=4.66, μd=-0.407, 95% CI=-0.530 to -0.284. For *TAS2R10*: (μ_normal_=2.52, μ_tumor_=1.92, μd=-0.603, 95% CI=-0.736 to -0.469. For *TAS2R14*: (μ_normal_=6.36, μ_tumor_=6.19, μd=- 0.168, 95% CI=-0.259 to -0.077. ***p<0.001; ****p<0.0001. **C)** Overall survival and **D)** progression-free survival Kaplan-Meier curve of 504 patients with PTC stratified by expression level of *TAS2R14* (high vs. low).

## Discussion

PTC is the most prevalent type of thyroid cancer and is rising in incidence.^41^ Resection with RAI therapy for high-risk pathology is the mainstay of treatment, but there are cases where this is either overtreatment with side-effects or inadequate treatment to achieve cure. For very low-risk PTC, there has been a paradigm shift towards less aggressive treatment such as thyroid lobectomy rather than total thyroidectomy, radiofrequency ablation (RFA), or active surveillance, but many healthcare providers are still hesitant to adopt surveillance as a management option.^9^ New therapeutic agents could bridge the gap between active surveillance and surgery and refine treatment further to balance effectiveness and morbidity. For aggressive subtypes of PTC, there is a risk of inadequate treatment due to a poor understanding of PTC tumor biology, making these tumors ripe for investigation of new therapeutic agents.^42^

T2R signaling is a promising, emerging field in cancer, and T2Rs may be new targets for non-invasive treatments for challenging cancers such as head and neck squamous cell carcinoma.^11,12^ While there has been encouraging work on T2Rs in other cancers, this is the first study to explore and characterize T2R expression and signaling in PTC cells. We show that T2R receptors regulate apoptosis in PTC cells, which provides a novel signaling pathway with potential clinical utility for PTC treatment, especially of low-risk or recurrent PTC.

This study is the first description of *TAS2R* expression in PTC cells. It was found that *TAS2R14* is the most consistently highly expressed *TAS2R* amongst the cell lines tested. This is in line with the expression level of *TAS2R14* in other cancers, such as head and neck squamous cell carcinoma.^11,14,43^ One of the PTC cell lines, MDA-T32 also had relatively high expression levels of several other *TAS2R*s including *TAS2R44*, *TAS2R45*, *TAS2R46*, and *TAS2R47*, possibly indicating a generalized upregulation of the taste receptor genes. The increased expression of *TAS2R*s in MDA-T32, a well-differentiated PTC, could correlate with the better prognosis of this variant compared to others, though this was not specifically explored. While this is the first study to find *TAS2R* expression in PTC cells, prior studies have found *TAS2R* expression in normal thyroid cells and tissue: 15 of 25 *TAS2R*s were analyzed via qPCR in thyrocytes and found to be expressed.^44^ In a mouse model, *TAS2R131* (a mouse TAS2R isoform most closely related to human TAS2R42) was found to be expressed in native thyrocytes but not in the parafollicular cells.^44,45^ In human cadaver thyroid tissue, *TAS2R38* was found to be expressed but localization could not be visualized via immunostaining.^44^ In terms of cellular localization in PTC cells, T2R14 appeared to be mostly intracellular through immunohistochemistry. This intracellular expression correlates with prior localization studies of T2R14 in other cancer cells.^7,10^ Intracellular expression of T2R14 in the endoplasmic reticulum (ER) would be consistent with previous literature regarding the location of GPCRs; however, future studies are needed to determine the subcellular T2R14 localization.^2,46–48^

In addition to expression, we found that bitter taste agonists significantly increase intracellular calcium levels through T2Rs in PTC cells, likely from the ER.^12,49–52^ All six bitter agonists elicited significant calcium responses in at least two cell lines. Pharmacologic experiments showed that inhibition of T2R14 resulted in a significantly reduced calcium response to agonists that activate T2R14, suggesting that at least some bitter agonists trigger a T2R14-dependent calcium release. Additional T2Rs may be activated by agonists that we tested that do not activate T2R14. Our findings are different from prior literature on T2R signaling in normal thyrocytes where bitter agonists (including denatonium benzoate, chloramphenicol, and colchicine) were not found to alter intracellular calcium in the absence of TSH. This prior work showed that T2R agonists *decreased* intracellular calcium in normal thyrocytes in the presence of TSH.^44^ This is a promising finding for the therapeutic potential of T2Rs in PTC because the cellular response to T2R activation differs between normal and cancerous thyroid cells.

However, further studies are needed to better characterize the specific differences and they may influence specificity of bitter therapies.

Bitter agonists induce downstream caspase activation, decreased viability, and increased apoptosis in PTC cells. Five bitter agonists (denatonium, diphenhydramine, quinine, thujone, and lidocaine) significantly decreased viability in all three cell lines. These agonists with efficacy include several FDA-approved drugs for other indications, potentially making them more readily available for repurposing in the future.^53–55^ FFA had variable responses, which could be due to the fact that FFA has known anti-inflammatory effects that may enhance proliferation through pathways that are independent of T2Rs. Diphenhydramine, thujone, and lidocaine consistently induced apoptosis in all three cell lines. These three agonists all signal through T2R14, consistent with previous literature in other types of cancers. Bitter agonists that activate T2R14 were found to increase caspase activation and Bax/Bcl-2 ratio in breast cancer^56,57^, bladder cancer^58,59^, cervical cancer^60^, gastric cancer^61,62^, and glioblastoma^63^. Previous literature has also proposed mechanisms of how T2Rs induce apoptosis. One possible mechanism is inhibiting proteasomal degradation, which causes accumulation of proteins meant for degradation, leading to downstream mitochondrial dysfunction and ROS.^12^ Another proposed mechanism could be T2R- activated intracellular and intranuclear calcium elevation, which leads to mitochondrial depolarization and apoptosis.^17,64–66^

TCGA analysis suggests that *TAS2R* expression alterations impact PTC or serve as genetic markers of changes that affect tumor biology and treatment efficacy. Specifically, higher *TAS2R14* expression is correlated with better progression free survival. Since T2Rs regulate apoptosis, tumors that overexpress T2Rs may have an improved prognosis because of a more robust regulation on proliferation or an *in vivo* response to endogenous bitter agonists in the tumor microbiome or intracellular metabolites. This theory also correlates with the overall lower expression of T2Rs in tumor cells compared to normal thyroid cells. Previous genetic studies have been conducted for PTC showing that genetic variations in certain *TAS2R*s modify papillary carcinoma risk.^67^ Individuals with *TAS2R*3/4 CC haplotype were found to be at a lower risk for

PTC than the remaining haplotypes. Our analysis did not find a correlation between *TAS2R4* expression and survival, but future analysis could stratify TCGA data by haplotype to further investigate. There was no significant association between *TAS2R* expression and overall survival, suggesting T2Rs may be associated with disease control but not necessarily longer survival.

Given the already high survival rates of PTC, focusing on disease control may be more valuable and impactful for patient quality of life. Therefore, given our analysis, it could potentially prove beneficial to use *TAS2R* expression as a risk-stratifying tool to help with treatment triaging.

Higher *TAS2R* tumor expression could be an assessment feature when considering less invasive treatments, such as higher surveillance.

Further elucidation of the T2R signaling pathways in PTC cells is warranted. This current study is promising for the development of potential therapeutic avenues for PTC. Unresectable or recurrent disease that has failed standard of care, may benefit from targeted treatment options such as percutaneous ethanol injections in select cases.^8^ Given the accessible location of the thyroid and cervical lymphatics and efficacy of ultrasound guided interventions, it may be feasible to deliver bitter agonists as local injections either independently or in conjunction with other therapies. Additionally, unresectable disease or suspected microscopic disease may respond to local irrigation of the tumor bed after surgical resection using bitterants like lidocaine which has well-established safety. Lidocaine, which activates T2R14, has been demonstrated to be effective in improving overall survival and disease-free survival when injected peritumorally in breast cancer patients undergoing surgical resection.^68^ If similar clinical responses are seen in PTC, then bitter therapies like lidocaine may be valuable in the treatment algorithm for aggressive recurrent PTC. In a similar vein, given the feasibility and practicality of injectable bitter agonists, low-risk PTC that border the threshold between active surveillance and surgery could benefit from the use of bitter agonists. It could lessen the burden of frequent medical management by allowing more time between ultrasound screens and provide improved peace of mind for both provider and patient. Further studies are needed to establish bitter agonists as therapeutic agents for PTC, but this preclinical study shows their promising potential in PTC management.

### Future Directions

While this current study establishes the presence of T2Rs and their role in apoptosis in PTC cells, substantial research is required to fully appreciate the role of T2Rs in PTC. Potential future directions include comparative studies with normal thyroid tissue to establish differences in *TAS2R* levels that may occur with malignant transformation. Additional animal studies or patient sample analyses in PTC may further enhance the development of T2R therapies in PTC.

## Conclusion

PTC cells express T2Rs primarily intracellularly, including T2R14 which has been studied in multiple cancers. Bitter agonists induce calcium responses through T2Rs with downstream caspase activation, decreased viability, and apoptosis. Higher *TAS2R14* expression also correlates with better progression-free survival. These results suggest that activation of T2Rs by bitter agonists may have therapeutic and risk-stratifying applications in PTC, warranting further characterization of the involved signaling pathways.

## Author Contributions

RMC and RJL conceptualized and visualized the study; KW, BH, ZAM, AM, and JT investigated and involved in formal analysis; KW wrote—original draft; KW, BH, RMC, and RJL wrote—review and editing; RMC and RJL involved in funding acquisition; RMC and RJL supervised the study.

## Acknowledgements

We thank Maureen Victoria (University of Pennsylvania) for technical assistance.

Presented and awarded Best Thyroid Poster at the AHNS 2024 Annual Meeting at COSM

---

**Supplemental Figure 1.**
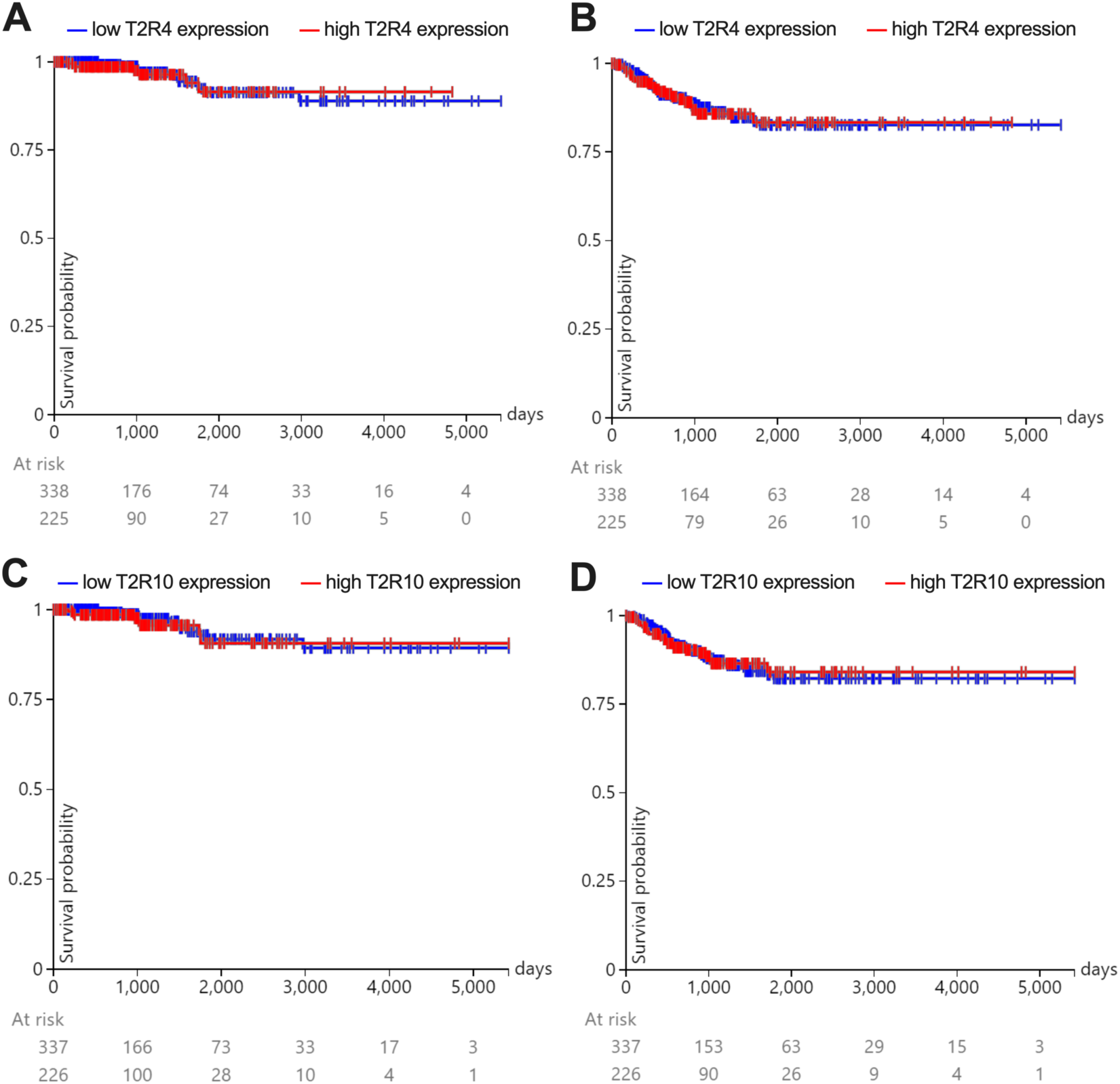
Kaplan-Meier curve of 504 patients with PTC. **A)** Overall survival plot and **B)** progression-free survival plot stratified by expression level of *TAS2R4* (high vs. low). **C)** Overall survival plot and **D)** progression-free survival plot stratified by expression level of *TASR10* (high vs. low).

## References

1. Pacini F, Castagna MG. Approach to and treatment of differentiated thyroid carcinoma. Med Clin North Am. 2012;96(2):369–383.

2. Hundahl SA, Fleming ID, Fremgen AM, Menck HR. A National Cancer Data Base report on 53,856 cases of thyroid carcinoma treated in the U.S., 1985-1995 [see commetns]. Cancer. 1998;83(12):2638-2648.

3. Noguchi S, Murakami N, Yamashita H, Toda M, Kawamoto H. Papillary thyroid carcinoma: modified radical neck dissection improves prognosis. Arch Surg. 1998;133(3):276–280.

4. Ito Y, Miyauchi A. Lateral and mediastinal lymph node dissection in differentiated thyroid carcinoma: indications, benefits, and risks. World J Surg. 2007;31(5):905–915.

5. Palme CE, Waseem Z, Raza SN, Eski S, Walfish P, Freeman JL. Management and outcome of recurrent well-differentiated thyroid carcinoma. Arch Otolaryngol Head Neck Surg. 2004;130(7):819–824.

6. Liu FH, Kuo SF, Hsueh C, Chao TC, Lin JD. Postoperative recurrence of papillary thyroid carcinoma with lymph node metastasis. J Surg Oncol. 2015;112(2):149–154.

7. Haddad RI, Bischoff L, Ball D, et al. Thyroid Carcinoma, Version 2.2022, NCCN Clinical Practice Guidelines in Oncology. J Natl Compr Canc Netw. 2022;20(8):925–951.

8. Goyal RM, Jonklaas J, Burman KD. Management of recurrent cervical papillary thyroid cancer. Endocrinol Metab Clin North Am. 2014;43(2):565–572.

9. Sugitani I, Ito Y, Takeuchi D, et al. Indications and Strategy for Active Surveillance of Adult Low-Risk Papillary Thyroid Microcarcinoma: Consensus Statements from the Japan Association of Endocrine Surgery Task Force on Management for Papillary Thyroid Microcarcinoma. Thyroid. 2021;31(2):183–192.

10. Singh N, Chakraborty R, Bhullar RP, Chelikani P. Differential expression of bitter taste receptors in non-cancerous breast epithelial and breast cancer cells. Biochem Biophys Res Commun. 2014;446(2):499–503.

11. Carey RM, McMahon DB, Miller ZA, et al. T2R bitter taste receptors regulate apoptosis and may be associated with survival in head and neck squamous cell carcinoma. Mol Oncol. 2022;16(7):1474–1492.

12. Miller ZA, Mueller A, Kim T, et al. Lidocaine induces apoptosis in head and neck squamous cell carcinoma through activation of bitter taste receptor T2R14. Cell Rep. 2023:113437.

13. Tuzim K, Korolczuk A. An update on extra-oral bitter taste receptors. J Transl Med. 2021;19(1):440.

14. Martin LTP, Nachtigal MW, Selman T, et al. Bitter taste receptors are expressed in human epithelial ovarian and prostate cancers cells and noscapine stimulation impacts cell survival. Mol Cell Biochem. 2019;454(1-2):203–214.

15. Freund JR, Mansfield CJ, Doghramji LJ, et al. Activation of airway epithelial bitter taste receptors by Pseudomonas aeruginosa quinolones modulates calcium, cyclic-AMP, and nitric oxide signaling. J Biol Chem. 2018;293(25):9824–9840.

16. Ozeck M, Brust P, Xu H, Servant G. Receptors for bitter, sweet and umami taste couple to inhibitory G protein signaling pathways. Eur J Pharmacol. 2004;489(3):139–149.

17. McMahon DB, Kuek LE, Johnson ME, et al. The bitter end: T2R bitter receptor agonists elevate nuclear calcium and induce apoptosis in non-ciliated airway epithelial cells. Cell Calcium. 2022;101:102499.

18. Pydi SP, Bhullar RP, Chelikani P. Constitutive activity of bitter taste receptors (T2Rs). Adv Pharmacol. 2014;70:303–326.

19. Goricanec D, Stehle R, Egloff P, et al. Conformational dynamics of a G-protein alpha subunit is tightly regulated by nucleotide binding. Proc Natl Acad Sci U S A. 2016;113(26):E3629–3638.

20. Wooding SP, Ramirez VA, Behrens M. Bitter taste receptors: Genes, evolution and health. Evol Med Public Health. 2021;9(1):431–447.

21. McMahon DB, Carey RM, Kohanski MA, Adappa ND, Palmer JN, Lee RJ. PAR-2- activated secretion by airway gland serous cells: role for CFTR and inhibition by Pseudomonas aeruginosa. Am J Physiol Lung Cell Mol Physiol. 2021;320(5):L845–L879.

22. McMahon DB, Carey RM, Kohanski MA, et al. Neuropeptide regulation of secretion and inflammation in human airway gland serous cells. Eur Respir J. 2020;55(4).

23. McMahon DB, Workman AD, Kohanski MA, et al. Protease-activated receptor 2 activates airway apical membrane chloride permeability and increases ciliary beating. FASEB J. 2018;32(1):155–167.

24. Sriram K, Moyung K, Corriden R, Carter H, Insel PA. GPCRs show widespread differential mRNA expression and frequent mutation and copy number variation in solid tumors. PLoS Biol. 2019;17(11):e3000434.

25. Carey RM, Kim T, Cohen NA, Lee RJ, Nead KT. Impact of sweet, umami, and bitter taste receptor (TAS1R and TAS2R) genomic and expression alterations in solid tumors on survival. Sci Rep. 2022;12(1):8937.

26. Gaida MM, Mayer C, Dapunt U, et al. Expression of the bitter receptor T2R38 in pancreatic cancer: localization in lipid droplets and activation by a bacteria-derived quorum-sensing molecule. Oncotarget. 2016;7(11):12623–12632.

27. Stern L, Giese N, Hackert T, et al. Overcoming chemoresistance in pancreatic cancer cells: role of the bitter taste receptor T2R10. J Cancer. 2018;9(4):711–725.

28. Banerjee P, Preissner R. BitterSweetForest: A Random Forest Based Binary Classifier to Predict Bitterness and Sweetness of Chemical Compounds. Front Chem. 2018;6:93.

29. Costa AR, Duarte AC, Costa-Brito AR, Goncalves I, Santos CRA. Bitter taste signaling in cancer. Life Sci. 2023;315:121363.

30. Chua SL, See Too WC, Khoo BY, Few LL. UBC and YWHAZ as suitable reference genes for accurate normalisation of gene expression using MCF7, HCT116 and HepG2 cell lines. Cytotechnology. 2011;63(6):645-654.

31. Christensen JN, Schmidt H, Steiniche T, Madsen M. Identification of robust reference genes for studies of gene expression in FFPE melanoma samples and melanoma cell lines. Melanoma Res. 2020;30(1):26–38.

32. Edelstein AD, Tsuchida MA, Amodaj N, Pinkard H, Vale RD, Stuurman N. Advanced methods of microscope control using muManager software. J Biol Methods. 2014;1(2).

33. Fierro F, Peri L, Hubner H, et al. Inhibiting a promiscuous GPCR: iterative discovery of bitter taste receptor ligands. Cell Mol Life Sci. 2023;80(4):114.

34. Goldman MJ, Craft B, Hastie M, et al. Visualizing and interpreting cancer genomics data via the Xena platform. Nat Biotechnol. 2020;38(6):675–678.

35. Spinelli KJ, Gillespie PG. Monitoring intracellular calcium ion dynamics in hair cell populations with Fluo-4 AM. PLoS One. 2012;7(12):e51874.

36. Kim D, Pauer SH, Yong HM, An SS, Liggett SB. beta2-Adrenergic Receptors Chaperone Trapped Bitter Taste Receptor 14 to the Cell Surface as a Heterodimer and Exert Unidirectional Desensitization of Taste Receptor Function. J Biol Chem. 2016;291(34):17616–17628.

37. Choudhary GS, Al-Harbi S, Almasan A. Caspase-3 activation is a critical determinant of genotoxic stress-induced apoptosis. Methods Mol Biol. 2015;1219:1–9.

38. Lamkanfi M, Kanneganti TD. Caspase-7: a protease involved in apoptosis and inflammation. Int J Biochem Cell Biol. 2010;42(1):21–24.

39. Brentnall M, Rodriguez-Menocal L, De Guevara RL, Cepero E, Boise LH. Caspase-9, caspase-3 and caspase-7 have distinct roles during intrinsic apoptosis. BMC Cell Biol. 2013;14:32.

40. Wei K, et al. Supplemental data to: Bitter Taste Receptor Agonists Induce Apoptosis in Papillary Thyroid Cancer. Figshare. 2024.

41. Lim H, Devesa SS, Sosa JA, Check D, Kitahara CM. Trends in Thyroid Cancer Incidence and Mortality in the United States, 1974-2013. JAMA. 2017;317(13):1338-1348.

42. Coca-Pelaz A, Shah JP, Hernandez-Prera JC, et al. Papillary Thyroid Cancer-Aggressive Variants and Impact on Management: A Narrative Review. Adv Ther. 2020;37(7):3112–3128.

43. Singh N, Shaik FA, Myal Y, Chelikani P. Chemosensory bitter taste receptors T2R4 and T2R14 activation attenuates proliferation and migration of breast cancer cells. Mol Cell Biochem. 2020;465(1-2):199–214.

44. Clark AA, Dotson CD, Elson AE, et al. TAS2R bitter taste receptors regulate thyroid function. FASEB J. 2015;29(1):164–172.

45. Lossow K, Hubner S, Roudnitzky N, et al. Comprehensive Analysis of Mouse Bitter Taste Receptors Reveals Different Molecular Receptive Ranges for Orthologous Receptors in Mice and Humans. J Biol Chem. 2016;291(29):15358–15377.

46. Crilly SE, Puthenveedu MA. Compartmentalized GPCR Signaling from Intracellular Membranes. J Membr Biol. 2021;254(3):259–271.

47. Jong YI, Harmon SK, O’Malley KL. GPCR signalling from within the cell. Br J Pharmacol. 2018;175(21):4026–4035.

48. Ribeiro-Oliveira R, Vojtek M, Goncalves-Monteiro S, et al. Nuclear G-protein-coupled receptors as putative novel pharmacological targets. Drug Discov Today. 2019;24(11):2192–2201.

49. McDonough RC, Gilbert RM, Gleghorn JP, Price C. Targeted Gq-GPCR activation drives ER-dependent calcium oscillations in chondrocytes. Cell Calcium. 2021;94:102363.

50. Wang Y, Sherrard A, Zhao B, et al. GPCR-induced calcium transients trigger nuclear actin assembly for chromatin dynamics. Nat Commun. 2019;10(1):5271.

51. Chakraborty S, Hasan G. Store-Operated Ca(2+) Entry in Drosophila Primary Neuronal Cultures. Methods Mol Biol. 2018;1843:125–136.

52. Gruntovskii G. [Ceramic plastic repair in treating giant cell bone tumors]. Ortop Travmatol Protez. 1986(8):4–5.

53. Sessler DI. Long-term consequences of anesthetic management. Anesthesiology. 2009;111(1):1–4.

54. Achan J, Talisuna AO, Erhart A, et al. Quinine, an old anti-malarial drug in a modern world: role in the treatment of malaria. Malar J. 2011;10:144.

55. Rosenberg MH, Blumenthal LS. The clinical uses of intravenous diphenhydramine hydrochloride. Am J Med Sci. 1948;216(2):158–162.

56. Moradzadeh M, Hosseini A, Erfanian S, Rezaei H. Epigallocatechin-3-gallate promotes apoptosis in human breast cancer T47D cells through down-regulation of PI3K/AKT and Telomerase. Pharmacol Rep. 2017;69(5):924–928.

57. Hallman K, Aleck K, Quigley M, et al. The regulation of steroid receptors by epigallocatechin-3-gallate in breast cancer cells. Breast Cancer (Dove Med Press). 2017;9:365-373.

58. Luo KW, Wei C, Lung WY, et al. EGCG inhibited bladder cancer SW780 cell proliferation and migration both in vitro and in vivo via down-regulation of NF-kappaB and MMP-9. J Nutr Biochem. 2017;41:56–64.

59. Park C, Cha HJ, Lee H, et al. Induction of G2/M Cell Cycle Arrest and Apoptosis by Genistein in Human Bladder Cancer T24 Cells through Inhibition of the ROS-Dependent PI3k/Akt Signal Transduction Pathway. Antioxidants (Basel). 2019;8(9).

60. Li L, Qiu RL, Lin Y, et al. Resveratrol suppresses human cervical carcinoma cell proliferation and elevates apoptosis via the mitochondrial and p53 signaling pathways. Oncol Lett. 2018;15(6):9845–9851.

61. Lu X, Li Y, Li X, Aisa HA. Luteolin induces apoptosis in vitro through suppressing the MAPK and PI3K signaling pathways in gastric cancer. Oncol Lett. 2017;14(2):1993–2000.

62. Li H, Lu H, Lv M, Wang Q, Sun Y. Parthenolide facilitates apoptosis and reverses drug-resistance of human gastric carcinoma cells by inhibiting the STAT3 signaling pathway. Oncol Lett. 2018;15(3):3572–3579.

63. Taylor MA, Khathayer F, Ray SK. Quercetin and Sodium Butyrate Synergistically Increase Apoptosis in Rat C6 and Human T98G Glioblastoma Cells Through Inhibition of Autophagy. Neurochem Res. 2019;44(7):1715–1725.

64. Gerasimenko OV, Gerasimenko JV, Tepikin AV, Petersen OH. Calcium transport pathways in the nucleus. Pflugers Arch. 1996;432(1):1–6.

65. Martelli AM, Mazzotti G, Capitani S. Nuclear protein kinase C isoforms and apoptosis. Eur J Histochem. 2004;48(1):89–94.

66. Petersen OH, Gerasimenko OV, Gerasimenko JV, Mogami H, Tepikin AV. The calcium store in the nuclear envelope. Cell Calcium. 1998;23(2-3):87–90.

67. Choi JH, Lee J, Yang S, Lee EK, Hwangbo Y, Kim J. Genetic variations in TAS2R3 and TAS2R4 bitterness receptors modify papillary carcinoma risk and thyroid function in Korean females. Sci Rep. 2018;8(1):15004.

68. Badwe RA, Parmar V, Nair N, et al. Effect of Peritumoral Infiltration of Local Anesthetic Before Surgery on Survival in Early Breast Cancer. J Clin Oncol. 2023;41(18):3318–3328.

69. Dagan-Wiener A, Di Pizio A, Nissim I, et al. BitterDB: taste ligands and receptors database in 2019. Nucleic Acids Res. 2019;47(D1):D1179–D1185.

